# Atypical BlaIR Two-Component System in *Pseudomonas aeruginosa* Regulates Virulence but not β-Lactam Resistance

**DOI:** 10.64898/2026.08.03.742534

**Authors:** Jonathan Ho, Wing Yin Venus Lau, Mila E. Tkatchouk, Michael J. Trimble, Manjeet Bains, Olga Pacios Santamaria, Alexandra Redey, Chloe Chan, Travis M. Blimkie, Negin Ketabchi, Patrick K. Taylor, Mahta Amanian, William W.L. Hsiao, Fiona S.L. Brinkman, Amy HY. Lee

**Affiliations:** Department of Molecular Biology and Biochemistry, Simon Fraser University, Burnaby, Canada; Faculty of Health Sciences, Simon Fraser University, Burnaby, BC, Canada; Department of Microbiology and Immunology, Centre for Microbial Diseases and Immunity Research, UBC, Vancouver, Canada; Department of Biochemistry and Molecular Biology, University of British Columbia

## Abstract

With the rise of antimicrobial resistance, anti-virulence therapeutics are a viable alternative to circumvent resistance pressures. Hypothetical genes and proteins are an under-studied source of potential virulence factor targets. We performed bioinformatic analyses to identify conserved hypothetical genes enriched in pathogenic *Pseudomonas aeruginosa* but not in non-pathogenic strains. This analysis identified an atypical BlaIR system, which we named *pvmSR,* that regulated *P. aeruginosa* virulence in a *Caenorhabditis elegans* infection model. This is in contrast with the typical BlaIR system from *Staphylococcus aureus*, which regulates resistance to β-lactam antibiotics. The *ΔpvmSR* mutant showed reduced virulence in a *C. elegans* slow-killing assay. To understand how PvmSR regulated virulence *in vivo*, we performed dual RNA-seq to analyze transcriptomic changes in both *C. elegans* and *P. aeruginosa.* We found that *C. elegans* responded to *P. aeruginosa ΔpvmSR* infection by decreasing expression of lysosome and phagocytosis pathways. In *P. aeruginosa ΔpvmSR,* we observed decreased gene expression of several known virulence factors including the hydrogen cyanide synthase, *hcnC*, and heparinase, *hepP*. Additionally, we observed dysregulation in genes important for quorum sensing and biofilm formation. Collectively, our findings indicated that PvmSR contributed to virulence regulation and may serve as a potential anti-virulence target.

## Introduction

In 2015, the World Health Organization (WHO) implemented the Global Action Plan (1) on Antimicrobial Resistance urging researchers to develop novel therapeutics and preventatives to combat antimicrobial resistance (2). Specifically, WHO established the following three criteria for drug research and development efforts towards alternative antimicrobial strategies to focus on therapeutics that can: (i) reduce pathogen virulence or host damage; (ii) minimize resistance development; and (iii) improve target specificity. Traditional antimicrobials often target essential proteins or processes and thus increase selective pressures on the bacterium leading to antimicrobial resistance (3). In contrast, anti-virulence therapeutics target virulence without killing the microbe and can be an attractive strategy in combating infections by increasing target specificity while minimizing resistance; specifically, anti-virulence therapeutics target proteins and toxins produced by pathogens during active infections and do not impact commensal bacteria present in the microbiome (3). By targeting virulence mechanisms, this also prevents host damage and allows the immune system time to mount an appropriate response with less morbidity (3).

One of the first steps in developing anti-virulence therapeutics is to identify potential virulence factors. Despite a high number of available sequenced bacterial genomes, at least one-third of predicted proteins lack clear functional annotation, due primarily to a lack of similarity to experimentally characterized proteins currently in public databases (4–7). In addition, predictions that are solely based on sequence similarities to experimentally characterized proteins can be incorrect due to functional divergence of homologous proteins (8). Thus, while there is a massive increase in available bacterial genome sequences, a large sequence-to-function knowledge gap impedes our abilities to functionally annotate predicted proteins from bacterial genomes (9, 10).

Previously, our group developed a bioinformatic approach to identify hypothetical proteins that are disproportionately found in annotated pathogens vs non-pathogens, named pathogen-associated genes (PAGs)(11). We hypothesize that such pathogen-associated hypothetical proteins, can serve as an untapped resource for the identification of potential novel virulence factors – and new targets for anti-virulence interventions. We previously characterized each gene from 298 pathogenic and 333 non-pathogenic bacterial genomes obtained from the National Center for Biotechnology Information (NCBI) RefSeq database, identifying those that were pathogen-associated, non-pathogen-associated, or “common” (found in both pathogens and non-pathogens)(11). We report here a refined version of this analysis and used this as a basis for further characterization of select WHO priority pathogens (12). Focusing on the priority pathogen *Pseudomonas aeruginosa*, this analysis identified an atypical BlaIR system that regulates *P. aeruginosa* virulence, using an established *Caenorhabditis elegans* virulence assay. In contrast to the canonical BlaIR two-component system found in *Staphylococcus aureus* (13, 14), which regulates resistance to β-lactam antibiotics, the BlaIR in *P. aeruginosa* did not play a role in regulating antibiotic resistance. Instead, deletion of this atypical BlaIR in *P. aeruginosa* reduced virulence in the *C. elegans* model, which we have renamed as PvmSR (*<u>P</u>seudomonas* <u>v</u>irulence/<u>m</u>etabolism <u>S</u>ensor and <u>R</u>egulator). To understand how PvmSR regulates virulence *in vivo*, we performed dual RNA-seq to assess both the host and bacterial responses at the same time. Our work showed that the *P. aeruginosa* PvmSR regulates iron sequestration, quorum sensing, and virulence genes in *P. aeruginosa* during infection, while altering *C. elegans* host responses to bacterial invasion. Collectively, our work illustrates the benefit of bioinformatically identifying pathogen-associated genes followed by functional characterization as a critical step towards anti-virulence candidate identification.

## Results

### Pathogen-associated genes analysis includes an atypical BlaIR system in *P. aeruginosa*

Previously, a comparative genomic analysis for identifying PAGs from comprehensive bacterial genome datasets was developed with the premise established by molecular Koch’s postulate that defined pathogen-associated genes as those genes exclusively conserved in pathogenic bacteria and absent in non-pathogenic bacteria (11). As newly sequenced bacterial genomes continue to increase, we updated the pathogen-associated genes analysis using 8,449 bacterial RefSeq genomes (downloaded from NCBI in 2018). From these RefSeq genomes, 328,750 (5.8%) patho-gen-associated genes, 837,103 (14.8%) non-pathogen-associated genes and 4,490,387 (79.4%) common genes were identified (**Figure S1**). Subsequently, we performed OrthoFinder-based orthology inference to distinguish orthologs from paralogs and improve the confidence in PAG identification (15). We then performed a genus-specific PAG analysis using 605 curated *Pseudomonas* genomes (**Figure 1**). As we were interested in experimentally assessing these PAGs, we prioritized PAGs with orthologs in more than three *Pseudomonas* pathogen genomes, with a focus on identifying novel candidates (i.e., hypothetical proteins without functional annotations) found in the virulent *P. aeruginosa* strain PA14. Furthermore, we prioritized candidates that also had evidence of positive selection to identify evolutionarily conserved putative virulence factors that may confer a benefit. While the degree to which a gene may be pathogen-associated may change, depending on the context, including genomes sequenced, positive selection will be maintained and is evidence of increased selective functional utility of a gene.

**Figure 1.**
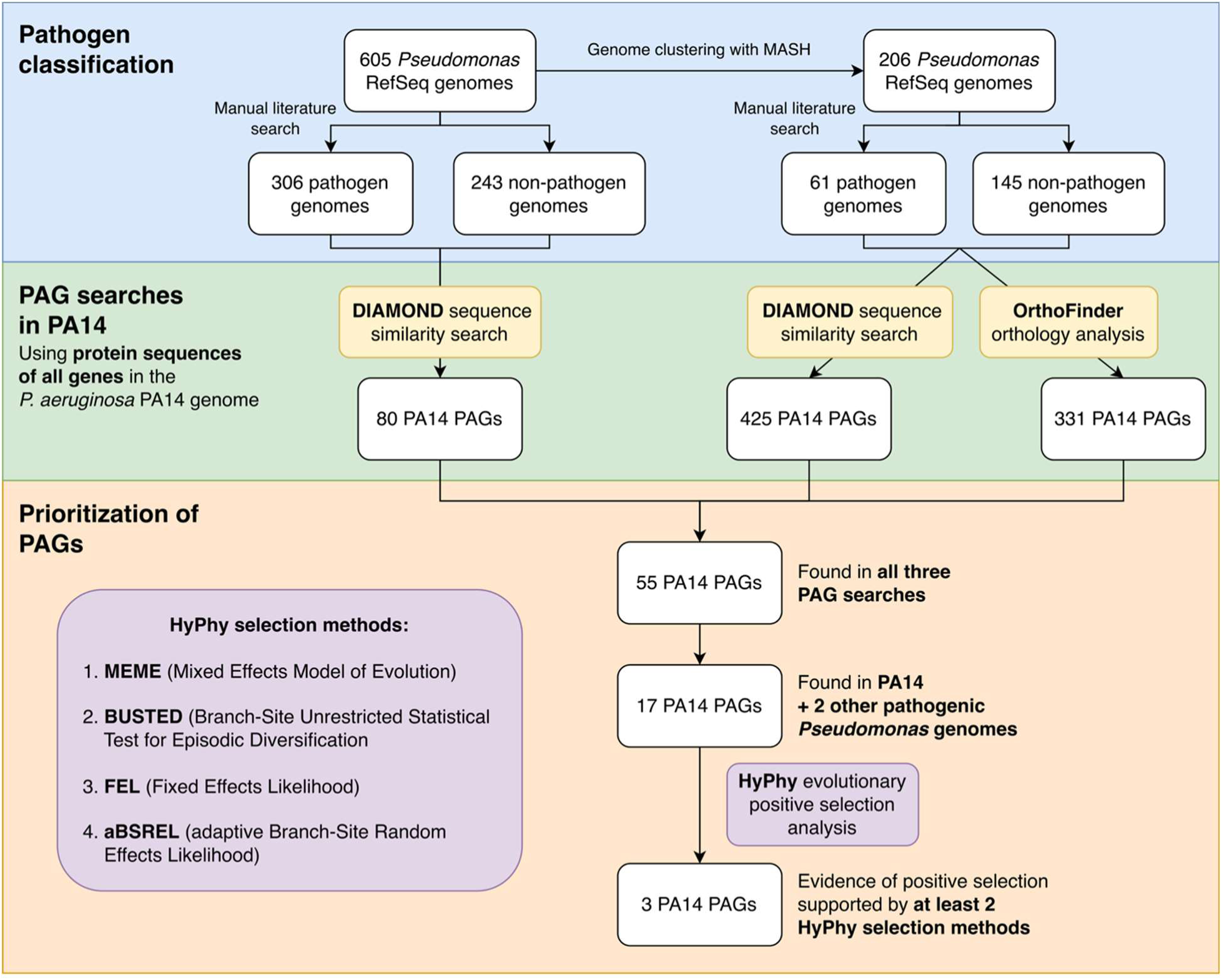
Workflow detailing the comparative bacterial genomic analysis used to identify PAGs in *P. aeruginosa* PA14. A complete genome set of 605 *Pseudomonas* genomes was obtained from RefSeq on October 12, 2020, from which a reduced set was obtained through genome clustering with MASH(55). Genomes were classified as pathogen or non-pathogen genomes through a manual literature search. Three searches were then done using the entire set of PA14 protein sequences; (1) a DIAMOND-based sequence similarity search against the complete genome set, (2) a DIAMOND-based sequence similarity search against the reduced genome set, and (3) an orthology analysis with OrthoFinder against the reduced genome set. PAGs were identified as being present only in pathogenic genomes, and PAGs found in all three searches, as well as being found in PA14 and at least two other pathogenic *Pseudomonas* genomes were prioritized for evolutionary positive selection with HyPhy (20).

Guided by the prioritization criteria above, we identified a two-component regulatory system consisting of the genes PA14_31050 (PA14_RS12695; WP_016254216.1) and PA14_31060 (PA14_RS12700; WP_025297936.1). Specifically, *Pseudomonas* Genome Database (PGDB) (16, 17) annotations for both PA14_31050 and PA14_31060 labeled them as hypothetical proteins. BLASTp analyses identified PA14_31050 as a M56 family metallopeptidase (90.33% identity, E ≈ 0), and PA14_31060 as a putative BlaI/MecI/CopY family transcriptional regulator (89.38% identity, E ≈ 0). PA14_31050, which we have named PvmS *(Pseudomonas* virulence/metabolism Sensor), was predicted to be under episodic positive selection (detected in some but not all branches of the gene tree) both at the site and gene-level based on combined results from MEME and BUSTED, respectively (**Table 1**)(18–20). The positive selection inference of this gene is further supported by aBSREL in which a proportion of branches on the gene tree were detected to be under positive selection(21). This metallopeptidase is predicted by PSORTb 3.0 (22) to localize in the cytoplasmic membrane.

**Table 1.** PAG analysis identified a potential BlaI/R transcriptional regulator and receptor. A subset of genes identified by positive selection analysis. Mixed Effect Model of Evolution (MEME) was used for detecting site-specific episodic positive selection, Branch-Site Unrestricted Statistical Test for Episodic Diversification (BUSTED) identifies gene-wide episodic positive selection, the Fixed Effects Likelihood (FEL) tests for pervasive positive selection, and the Adaptive Branch Site Random Effects Likelihood (aBSREL) identifies specific lineages that have clear positive selection. Putative functions for genes are based on BLAST prediction and NCBI’s prokaryotic genome annotation pipeline (PGAP). “Yes” denotes positive selection.

| Old locus tag | New locus tag | Gene name | Putative function | MEME | BUSTED | FEL | aBSREL |
| --- | --- | --- | --- | --- | --- | --- | --- |
| PA14_31050 | PA14_RS12695 | *pvmS* | M56 family metallopeptidase | Yes | Yes | No | Yes |
| PA14_31060 | PA14_RS12700 | *pvmR* | Blal/MecI/CopY transcriptional regulator | No | No | No | No |

Based on a multiple sequence alignment with other M56 family proteases using Clustal Omega and a transmembrane helix prediction using TMHMM Server 2.0 (**Figure S2A-B**) respectively(23, 24), PvmS possesses the M56 family signature HEXXH zinc-binding motif, between the third and fourth transmembrane domains, followed by an aspartate five residues downstream of the second histidine in the motif (25). Expression of PvmS is likely controlled by the transcriptional regulator PA14_31060, which is directly downstream; although this gene is not under positive selection (**Table 1**), its putative role as a regulator and predicted organization with PvmS on an operon (from PGDB;(16, 17)) warrants the investigation of these two genes as a system. We have therefore named PA14_31060, PvmR *(Pseudomonas* virulence/metabolism Regulator). Both the positively selected PvmS and its putative regulator, PvmR, are located on a genomic island based on IslandViewer 4 prediction, supported by Islander and IslandPick (**Figure S3**)(26).

Due to the genomic context of these two proteins and their annotations, we hypothesized that PvmSR in *P. aeruginosa* PA14 are functionally similar to other M56 proteins and their regulators, including the Bla and Mec systems in *S. aureus* involved in β-lactam resistance (27). Using AlphaFold 3 to generate a 3D structure of PvmS, we observed that PvmS was missing the cytoplasmic β-lactam sensor domain seen in the canonical BlaR1 from *S. aureus* (**Figure 2C, S4A**). A recent paper by Bonefont *et al.* (28) found a similar atypical BlaIR system in *Mycobacterioides abscessus* (formerly *Mycobacterium*), where the BlaR ortholog was also missing this sensor domain. Therefore, FoldMason multiple sequence alignments (MSAs) were done to compare the protein sequence of *P. aeruginosa* PvmSR with those of *S. aureus* BlaIR, *M. abscessus* and *M. tuberculosis* BlaIR from Bonefont *et al.*, and other orthologs from *Mycobacterium leprae* and *Nocardia brasiliensis* which were found in Foldseek structural similarity searches (**Figure 2A-B**). The PvmS MSA (**Figure 2A**) confirmed that, similarly to *M. abscessus* BlaR, PvmS contained the conserved HEXXH and EXXXD zinc ligand motifs and Arg294 of *S. aureus* BlaR1. In contrast, AlphaFold 3 prediction of PvmR showed that it has an additional N-terminal domain of poor confidence (1-77 aa, **Figure S4B**) compared to *S. aureus* BlaI (**Figure 2D**); however, the MSA of PvmR with the other BlaI homologs in **Figure 2A** revealed that no other protein had this domain.

**Figure 2.**
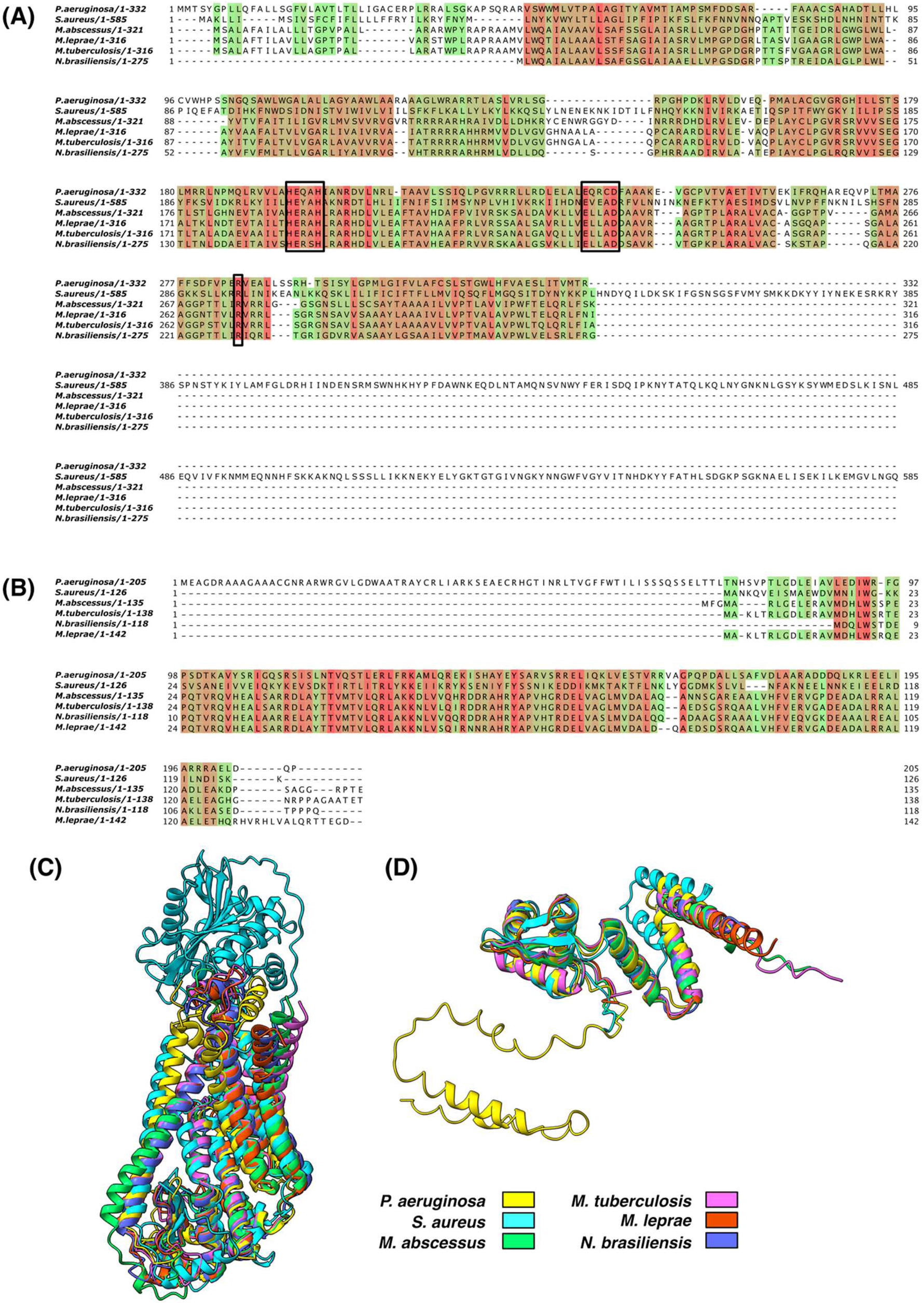
PvmSR are atypical homologs of BlaIR. **(A and B)** Protein sequence alignments of PvmS and PvmR to similar proteins from pathogenic species found in a Foldseek search in the AFDB-proteome database. Alignments were generated from a FoldMason structural alignment and visualized in Jalview (v2.11.5.1) (66). Alignments are coloured by conservation, with red being maximum and green being minimum. **(A)** PvmS aligned to *S. aureus* BlaR1 and atypical BlaR orthologs from other species. Key conserved residues from the BlaR1 active site (HEXXH and EXXXD motifs, stabilizing arginine) are marked in black boxes. PvmS is missing the periplasmic sensor domain, similar to atypical BlaR orthologs found in *M*. *tuberculosis* and *M. abscessus*. **(B)** PvmR aligned to *S. aureus* BlaI and atypical BlaI orthologs from other species. Sequence alignment indicates that PvmR has an additional N-terminal domain. **(C and D)** Structural alignments of PvmS **(C)** to the sequences in (A), and PvmR **(D)** to the sequences in **(B)**. 3D structures were generated using AlphaFold 3 (65) and visualized in ChimeraX (v1.11.1)(67). Structures are coloured as described in the legend.

### Deletion of pvmSR in *P. aeruginosa* reduces virulence in the *C. elegans* slow-killing assay

To assess the contributions of *pvmS* and *pvmR* to the virulence of PA14, we generated a marker-less deletion mutant *ΔpvmSR*. After confirmation of the deletion mutation using long-read sequencing (**Figure S5**), we evaluated the virulence of the deletion mutant *ΔpvmSR* using the established *C. elegans* slow-killing virulence assay (29). In contrast to the fast-killing assay where heat-killed bacteria can still mediate killing via toxins, slow-killing requires the colonization of the *C. elegans* gut by *P. aeruginosa*, which allows us to assess PvmSR’s potential virulence regulation. In brief, *P. aeruginosa* PA14 wildtype, PA14 *ΔpvmSR* or negative control *Escherichia coli* OP50 were grown on nematode growth medium (NGM) plates with 5-fluro-2ʹ-deoxyuridine (FUdR) overnight, followed by seeding 30 stage L4 worms, in triplicate, and monitored over eight days (**Figure 3A-B**). To assess the potential virulence impact of *pvmSR* mutations, *C. elegans* survival probability over eight days was analyzed using the Kaplan-Meier survival analyses. Across two biological replicate experiments (**Figure 3C** and **S6**), *ΔpvmSR* displayed decreased virulence as shown by the significantly lower hazard ratio compared to wildtype PA14 (HR = 0.46–0.61, q-value < 0.001; **Fig. S6B**).

**Figure 3.**
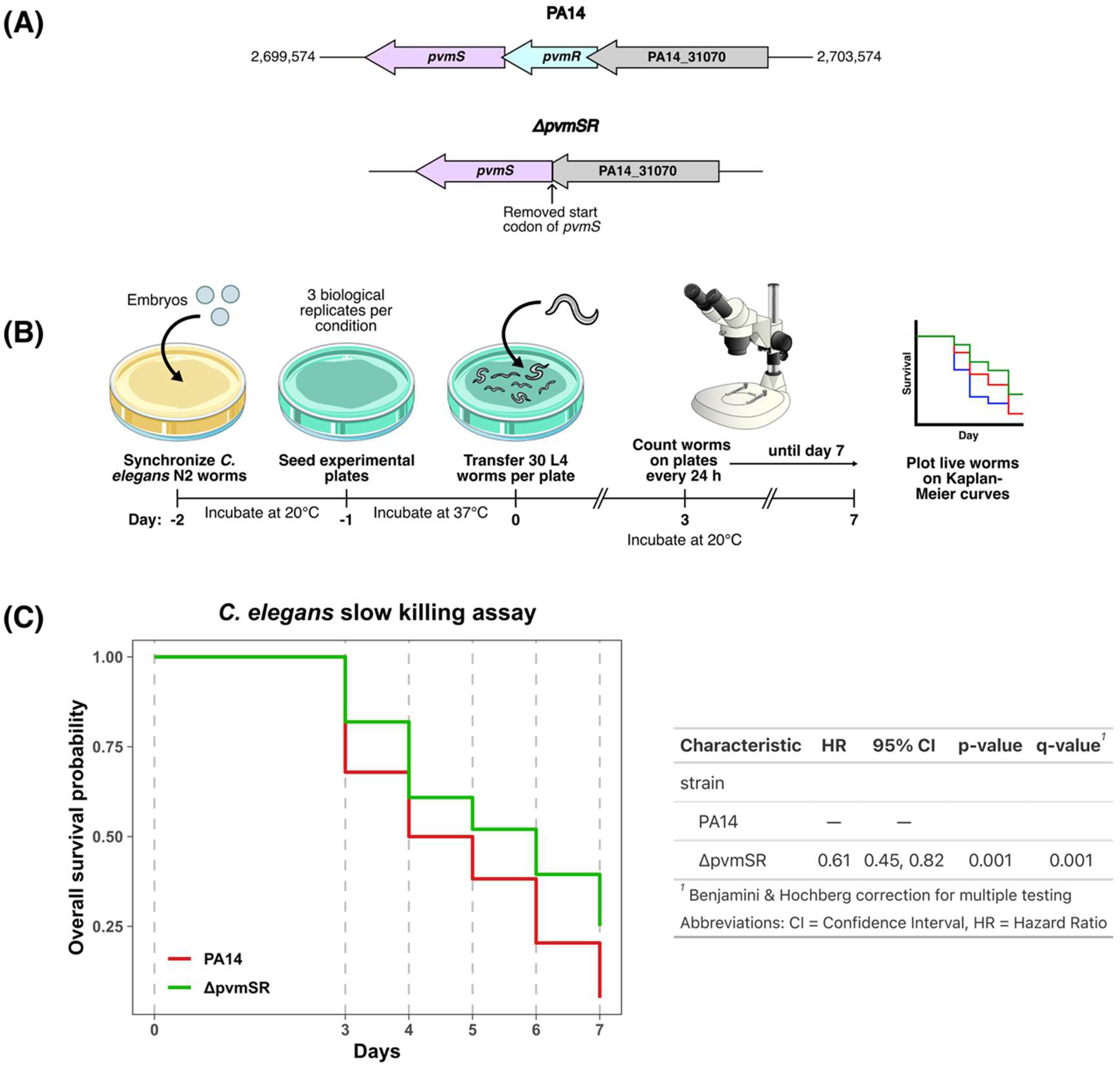
*C. elegans* slow killing assays comparing the virulence of the PvmSR mutant and wildtype PA14. **(A)** The genomic context of *pvmSR* in wildtype PA14 and in *ΔpvmSR*. *ΔpvmSR* was created through a clean deletion of *pvmR*, resulting in the deletion of the start codon of *pvmS*. **(B)** A timeline of the *C. elegans* slow-killing virulence assay. After synchronization, 30 L4 Bristol N2 *C. elegans* were placed onto plated bacterial lawns cultivated on NGM media supplemented with FUdR. Worm survival was monitored daily for 5 days and subsequently plotted on a Kaplan-Meier curve. **(C)** Kaplan-Meier curve of *C. elegans* slow-killing assay comparing the virulence of wildtype PA14 to *ΔpvmSR.* The graph is a representative of an experiment performed in triplicate. HR: hazard ratio, CI: confidence interval.

### *P. aeruginosa* PvmSR does not regulate β-lactam resistance

Given that PvmR showed protein sequence similarity to BlaI (21.70% identity, E = 7e-09), which regulates β-lactam resistance in *S. aureus*, we assessed β-lactam resistance of wildtype PA14 and *ΔpvmSR* using the following β-lactam antibiotics; ampicillin, carbenicillin, ceftazidime, and piperacillin. The mutant *ΔpvmSR* had the same minimum inhibitory concentrations (MICs) as seen in wildtype PA14 (**Table 2**). Additional classes of antibiotics were tested for increased resistance or sensitivity (nalidixic acid, kanamycin, and polymyxin B), but no MIC differences were observed (**Table 2**). Thus, unlike the BlaIR system in *S. aureus*, these *P. aeruginosa* homologs do not regulate antibiotic resistance.

**Table 2.** PA14 and the *ΔpvmSR* mutant have similar sensitivity to antibiotics in broth microdilution assay. Strains were cultured in MHB medium in the presence of serial dilutions of antibiotics to determine the minimum inhibitory concentration. The resulting analysis indicates that the *pvmSR* operon does not appear to affect antibiotic resistance to β-lactam, aminoglycoside, quinolone, and polymyxin antibiotics. MIC: minimal inhibitory concentration, pBBR1: pBBR1MCS-2 plasmid vector.

| | MIC (μg/mL) | | | | | | |
| --- | --- | --- | --- | --- | --- | --- | --- |
|  | Ampicillin | Carbenicillin | Polymyxin B | Ceftazidime | Piperacillin | Kanamycin | Nalidixic Acid |
| PA14/pBBR1 | 500 | 125 | 1.56 | 3.13 | 4 | >250 | 75 |
| $\Delta pvmSR$/pBBR1 | 500 | 125 | 1.56 | 3.13 | 4 | >250 | 75 |

### *C. elegans* infected with *P. aeruginosa ΔpvmSR* had down-regulated host responses to pathogen infection including phagocytosis and lysosome functions

To understand the bacterial and host transcriptomic changes that occur during PA14 colonization and infection of *C. elegans*, we performed dual RNA-seq on a modified protocol of the *C. elegans* slow-killing virulence assay as shown in **Figure 4A**. In brief, *P. aeruginosa* PA14 wildtype, and PA14 *ΔpvmSR* were grown on NGM plates with FUdR overnight, followed by seeding 100 stage L4 worms, in triplicates, and incubated over 24 hours. Infected worms were then processed for total RNA extraction for dual RNA-seq library preparation (see Methods). A median of 2.4M reads were uniquely mapped to both *C. elegans* and PA14.

**Figure 4.**
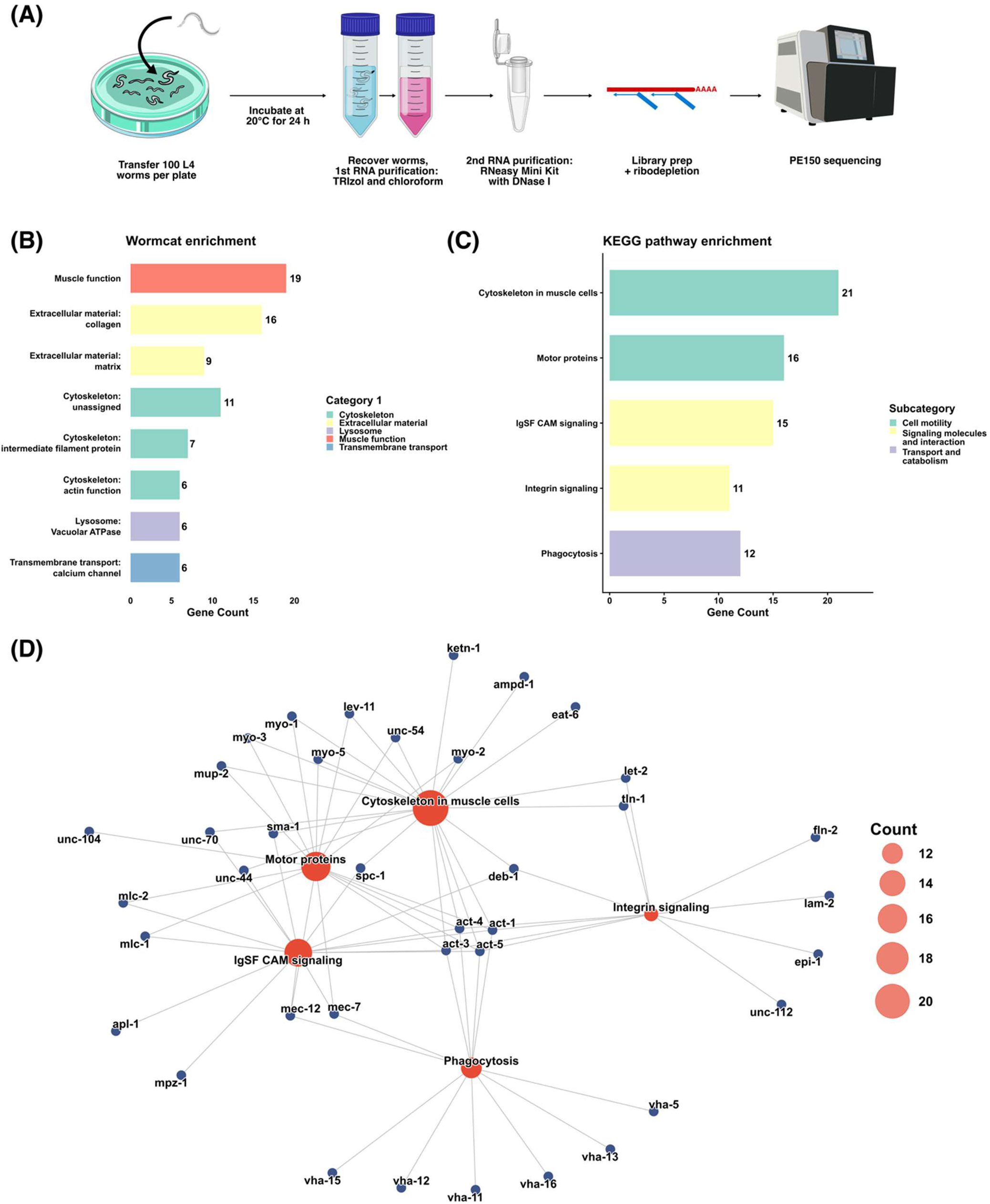
*C. elegans* transcriptomic response to infection by *ΔpvmSR* in contrast to wildtype PA14. **(A)** Schematic of the dual RNA-seq workflow to identify virulence changes in *ΔpvmSR* and host response in *C. elegans*. **(B)** Functional enrichment of Wormcat categories and **(C)** KEGG pathways in *C. elegans* infected by *ΔpvmSR* calculated using DEGs with an adjusted p-value ≤ 0.05 and fold change ≥ 1.5. Enriched descriptions are shown on the y-axis, with bars coloured by the broader category or pathway descriptions. Gene counts are shown and labelled with bars. **(D)** Gene-concept network of enriched KEGG pathways and their DEGs. Pathway nodes are coloured orange with size determined by gene counts. Gene nodes are coloured blue and connected to the pathways they enrich.

Principal component analysis showed that *C. elegans* samples separated by genotype of the infecting PA14 strain, with replicates clustering together on the first principal component representing 80% of the variance (**Figure S7A**). Differential expression analysis of the *C. elegans* host colonized by PA14 *ΔpvmSR* vs. PA14 wildtype identified 287 differentially expressed genes (DEGs) (21up, 266 down, using an adjusted *p*-value ≤ 0.05 and an absolute fold change ≥ 1.5; **Supplementary Table S1**; **Figure S7B**).

Several classes of genes were notably down-regulated: myosin, myosin regulatory light chain, cuticle collagen, actin, intermediate filament, and V-type proton ATPase genes (**Supplementary Tables S2** and **S3**). These genes drive functional enrichment in Wormcat categories consisting of muscle function, extracellular material, cytoskeleton, lysosome, and transmembrane transport (**Figure 4B** and **Supplementary Table S2**). Similar results were also found in enriched KEGG pathways, which included cytoskeleton in muscle cell, motor proteins, IgSF CAM signaling, integrin signaling, and phagocytosis (**Figures 4C**, **D** and **Supplementary Table S3**). These enriched pathways were interconnected through many of the aforementioned down-regulated genes, with the actin genes functioning across all the enriched pathways (**Figure 4D**). Taken together, decreased gene expression in muscle function, cytoskeleton, lysosome, and phagocytosis functions in *C. elegans* infected with PA14 *ΔpvmSR* compared to PA14 wildtype indicate a decreased host response to the PA14 *ΔpvmSR*.

### Deletion of *pvmSR* resulted in metabolism- and virulence-specific transcriptional remodelling in *P. aeruginosa* during infection

To understand how PvmSR regulates virulence during infection, we analyzed *in vivo* bacterial gene expression differences between PA14 *ΔpvmSR* versus wildtype during *C. elegans* infection using dual RNA-seq (**Figure 5A**). Principal component analysis of top 3000 most variable genes showed samples clustering by genotypes (**Figure S8A**), with negligible counts observed for both *pvmS* and *pvmR* gene in the *ΔpvmSR* mutant as expected (**Figure 5A**). Interestingly, we observed that in the PA14 wildtype, all the reads piled up at position 2,701,059 (219 nt downstream of the annotated start of *pvmR*), suggesting that this gene may be misannotated in the PA14 genome.

**Figure 5.**
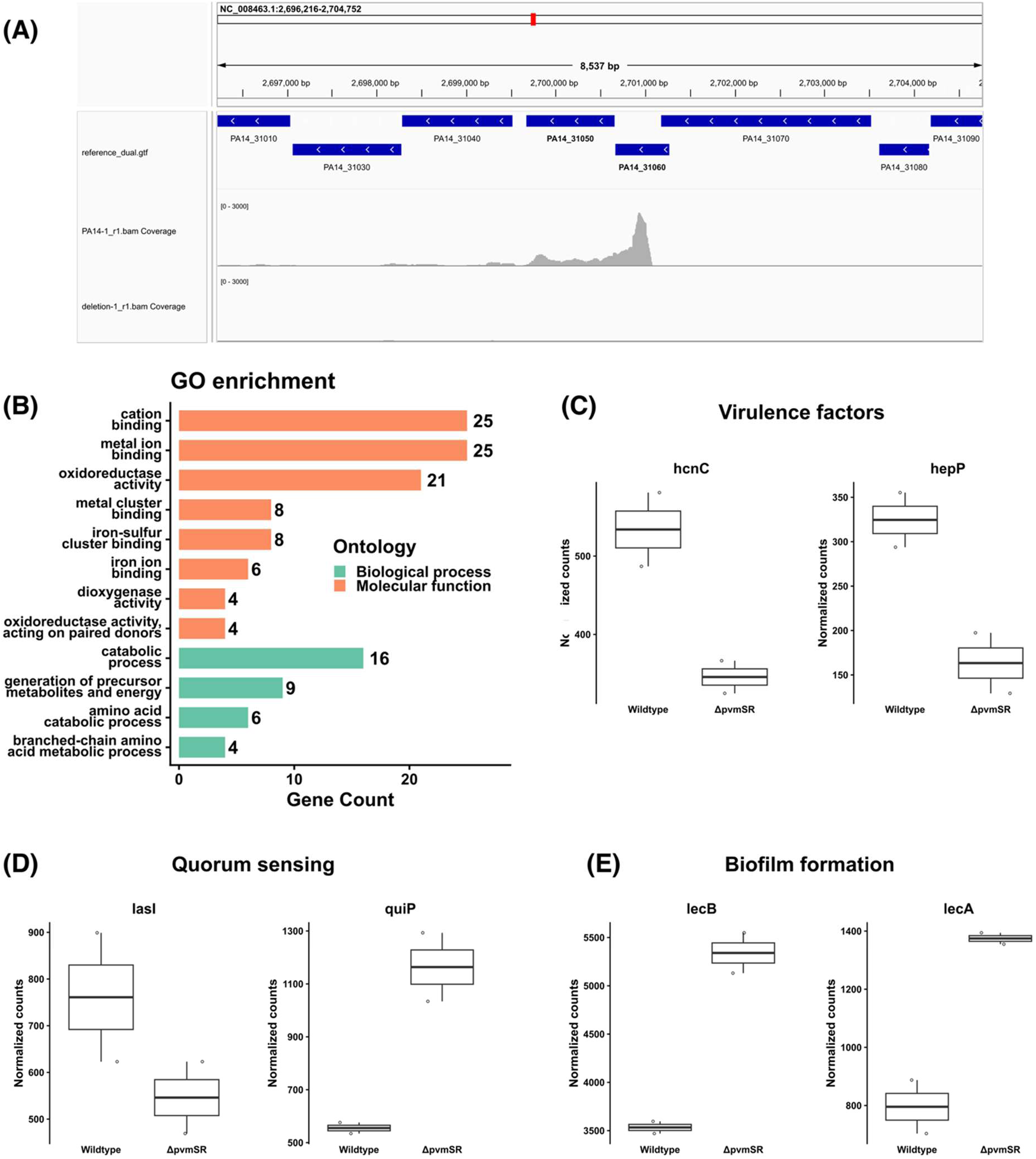
Transcriptomic profiles of *ΔpvmSR* in a modified *C. elegans* slow-killing virulence assay. **(A)** Read pileup of transcript sequences aligned to the *pvmSR* region in UCBPP-PA14_109 PA14 using Integrative Genomics Viewer (IGV). Genomic coordinates are shown on the top scale, and read coverage is represented as a density plot across wildtype and *ΔpvmSR* PA14. **(B)** Over-representation analysis of GO terms (adjusted p-value ≤ 0.05) in *ΔpvmSR* PA14 using all DEGs with an adjusted p-value ≤ 0.05 and fold change ≥ 0. Enriched term descriptions are shown on the y-axis, colours represent the ontologies the terms belong to, and gene counts are shown and labelled with bars. **(C-E)** DESeq2 normalized plot counts of genes associated with **(C)** virulence factors, **(D)** quorum sensing, and **(E)** biofilm formation for wildtype and *ΔpvmSR* PA14. Boxplots display medians with lower and upper hinges representing first and third quartiles; whiskers reach the highest and lowest values no more than 1.5× interquartile range from the hinge.

Differential expression analysis of the *ΔpvmSR* identified 121 DEGs (49 up, 72 down) with adjusted *p*-value of 0.05 (**Figure S8B, Supplementary Table S4)**. These DE genes were then mapped to potential Gene Ontology (GO) terms using eggnog-mapper, followed by GO enrichment analysis. Comparing *ΔpvmSR* to wildtype PA14 during infection, we observed a statistically significant enrichment in the dysregulation of GO biological processes that were related to amino acid metabolic processes, and the generation of precursor metabolites and energy (**Figure 5B, Supplementary Table S5**). Specifically, we observed enrichment of metabolic processes including oxidation-reduction reactions, aromatic and branched-chain amino acid metabolism, central carbon metabolism and respiratory electron transfer. Interestingly, the molecular functions of the DEGs showed a “GO Molecular Functions” enrichment of metal ion binding and iron-sulfur binding functions (**Figure 5B, Supplementary Table S5**).

To further understand how the *ΔpvmSR* mutant had decreased virulence in *C. elegans*, we examined the list of DEGs and identified dysregulation in virulence factors, quorum sensing and biofilm formation. These included the down-regulation of virulence factors including: the hydrogen cyanide biosynthesis gene *hcnC* (30), the heparinase *hepP* and its co-translated putative zinc-binding dehydrogenase *zbdP* (31), previously shown to be important for virulence in *C. elegans* (**Figure 5C**; **Supplementary Table S4**). We also observed down-regulation of *rsmA*, a global regulator of virulence-related genes important for initial colonization during infection (**Supplementary Table S4**) (32, 33).

Furthermore, we observed dysregulation in quorum sensing (QS) and biofilm in the *ΔpvmSR* mutant during infection. In *P. aeruginosa*, there are four QS systems: LasI/R, RhlI/R, Pqs quinolone system and the IQS system for phosphate-limiting conditions (34). Particularly, we observed a down-regulation of *lasI*, which produces the QS autoinducer acyl-homoserine lactone (AHL)(35); and an up-regulation of *quiP* (annotated as *pac* in the Pseudomonas Genome Database), the AHL acylase that degrades the QS autoinducer (36) (**Figure 5D**; **Supplementary Table S4**). We also observed dysregulation in various genes involved in biofilm formation in the *ΔpvmSR* mutant, including up-regulation in the anti-Sigma factor, MucA and genes encoding for lectins LecA and LecB (**Figure 5E**; **Supplementary Table S4**). Collectively, our dual RNA-Seq illustrated an uncoupling of key regulation to maintain successful infection when PvmSR is disrupted, leading to reduced expression of virulence factors and dysregulation in quorum sensing and biofilm regulation.

## Discussion

Anti-virulence therapeutic strategies hold promise in addressing antimicrobial resistance by targeting virulence factors to reduce pathogen virulence while minimizing impact on beneficial commensal bacteria and potentially reducing potential for resistance development. However, one of the major challenges in identifying putative anti-virulence candidates is the need for high-quality gene functional annotation, particularly for any novel, pathogen-specific genes that have limited sequence or structural similarity to known genes. Computational methods, including improved structural and sequence alignment models such as AlphaFold and FoldMason, are rapidly aiding the identification and functional assignments of hypothetical proteins (37–39). However, despite these technologies, accurate genome annotation remains challenging due to several factors; the presence of multiple potential start codons, short or overlapping open reading frames, and the limitations of automated gene prediction algorithms. Thus, the importance of refining these annotations using molecular approaches is therefore critical.

Using a comparative bacterial genomics approach (**Figure 1**), we identified a novel two-component regulatory system in *P. aeruginosa*, which we have named PvmSR *(Pseudomonas* virulence/metabolism sensor/regulator). Though sequence and structural alignments identified similarity to the BlaIR system in *S. aureus*, we found that PvmSR does not regulate β-lactam resistance (**Table 2**), highlighting the limits of homology-based functional annotation and the importance of experimental validation. This finding is supported by the lack of *S. aureus* BlaR’s canonical β-lactam sensor domain in PvmS (**Figure 2)**, raising questions as to what this regulatory system is sensing and responding to. The original *P. aeruginosa* PA14 genome, available in the PGDB (16, 17), annotated PvmR with an extended N-terminal domain, which is likely a misannotation. Specifically, this domain was modeled with low confidence in AlphaFold 3 and was not present in any other identified BlaI orthologs (**Figure 2**). Furthermore, in our dual RNA-seq experiments, we observed that transcription started 219 nt downstream from the annotated start codon, ansupporting that a change in annotation of the start codon is needed (**Figure 5**). Of note, the PA14 RefSeq annotation was updated in February 2025, providing a new annotation for *pvmR* (PA14_RS12700) that omits amino acid residues 1-73 (the first 219 nt).

Using an established *C. elegans* slow-killing virulence assay, we demonstrated that PvmSR is important for virulence, as PA14 *ΔpvmSR* showed reduced virulence compared to PA14 wildtype (**Figure 3**). To understand how the virulence of PA14 *ΔpvmSR* in *C. elegans* was reduced, we performed dual RNA-Seq to assess transcriptional changes in both the host and the pathogen (**Figure 4**). In *C. elegans* infected by *ΔpvmSR* compared to wildtype PA14, we observed decreased gene expression of V-type ATPases, with an overall functional enrichment of lysosomal and phagocytosis functions (**Figure 4**). In *C. elegans*, the acidification of lysosomes is driven by V-ATPase (40). Counterintuitively, mutations in V-ATPase that reduce V-ATPase levels and lead to dysfunction in lysosomes, which activate the innate immune system and promote pathogen resistance in *C. elegans* (40).

We next assess how PvmSR regulate *P. aeruginosa* virulence in the *C. elegans* host by analyzing pathogen-specific reads from dual RNA-Seq (**Figure 4**). Specifically, we saw that several known *C. elegans* virulence factors, including the cyanide biosynthesis gene *hcnC* and heparinase (*hepP*) were significantly down-regulated in the PA14 *ΔpvmSR* mutant (**Figure 5**)(31, 41). *hcnC* encodes a subunit of hydrogen cyanide synthase responsible for producing hydrogen cyanide; this toxin inhibits cytochrome C oxidases in host cells or competing microorganisms, and is the principal toxic factor responsible for rapid paralysis of *C. elegans* by *P. aeruginosa* (41). Although slow-killing mainly depends on intestinal colonization, reduced *hcnC* expression will likely lead to decreased cyanide production and reduced pathogenesis (41). Previous work has shown that PA14 grown in whole blood from severely burned patients up-regulates the expression of *hepP* and *zbdP* (31). HepP was shown experimentally to be a functional heparinase, and a transposon-insertion *hepP* mutant in PA14 showed reduction in pellicle and biofilm formation at the air-liquid interface, as well as reduced virulence in the *C. elegans* slow-killing model (31). Interestingly, a transposon-insertion *zbdP* mutant, which encodes a putative zinc-binding dehydrogenase showed increased biofilm formation (31), but conflicting *C. elegans* virulence phenotype (31, 32, 42), thus warranting further characterization.

Quorum sensing is a major regulator of *P. aeruginosa* virulence and biofilm formation(32). In the *ΔpvmSR* mutant, we observed dysregulation in two genes from the LasI/R QS system; specifically, the down-regulation of AHL synthase *lasI*, and the up-regulation of AHL acylase *quiP* (**Figure 5**). LasI synthesizes 3-oxo-C12-HSL(43), which binds and activates LasR to regulate the expression of numerous QS-controlled virulence genes(44). In contrast, QuiP cleaves long-chain AHL molecules and therefore functions as a quorum quenching enzyme(36). We hypothesize that the down-regulation of the AHL synthase gene *lasI* and the up-regulation of the AHL acylase gene, *quiP* likely leads to an overall reduction of the Las signal and quorum sensing communication. This could contribute to reduced virulence because enzymatic depletion of *P. aeruginosa* AHLs attenuates infection in *C. elegans*(45). Interestingly, the anthranilate degradation pathway encoded by *antABC* operon is also down-regulated, which may influence the *Pseudomonas* quinolone signal (PQS) QS system (46, 47) (**Supplementary Table S4**).

Another important component of virulence in *P. aeruginosa* is its ability to form biofilms, which facilitate persistent colonization and protect bacterial cells from host defenses and antibiotic treatment (48). We observed dysregulation in biofilm-associated genes in the *ΔpvmSR* mutant, including up-regulation of the anti-Sigma factor, MucA, and genes encoding for lectins, LecA and LecB (**Figure 5**). MucA is known to repress alginate production by repressing *algU* expression. In contrast, LecA and LecB are galactose- and fucose-binding lectins, respectively, that contribute to biofilm formation by promoting bacterial adhesion and stabilizing the biofilm matrix(49, 50). Together, these findings suggest that deletion of *pvmSR* leads to uncoordinated biofilm-associated gene expression during infection.

Collectively, this approach has identified an important set of sensor-regulator virulence genes, previously annotated as hypothetical genes, that may be suitable anti-virulence drug targets, and warrant further study as a potential master regulator system. This approach can be used to further identify other hypothetical genes that may play a role in virulence, that aren’t easily characterized by other means, due to the contextual nature of virulence phenotypes.

## Methods

### Comparative bacterial genomics identified pathogen-associated genes from the *Pseudomonas* genus

An updated set of 8646 bacterial genomes from the NCBI RefSeq was retrieved from the MicrobeDB database v97 (51) on April 26^th^, 2018. Genomes were assigned a pathogen status of “pathogen” or “non-pathogen” based on a manual curation of documented pathogenicity at the species level. Genomes were classified as a “pathogen” if their corresponding species had published evidence of infection in any host organism (e.g.: human, animal or plants), and were classified as “non-pathogens” if their corresponding species have been documented in publications as environmental organisms or as host-associated organisms with no current evidence of pathogenicity. Genomes that were not reported or characterized in any publications at the time of the curation were assigned an “unknown” pathogen status and were excluded from the analysis. All genomes/strains within a species were collectively assigned as a “pathogen” if at least one strain within the species is pathogenic in at least one host organism, or as a “non-pathogen” if none of the strains within the species is reportedly pathogenic towards any host organisms (11, 52).

Genes from the deduced proteomes of a total of 5196 pathogenic and 3253 non-pathogenic RefSeq genomes were searched for sequence similarity against all genes from all proteomes, excluding the proteome of origin, using DIAMOND(53, 54). An e-value cut-off of 10^-7^ was used to exclude potentially distant homologs. A gene is considered pathogen-associated, non-patho-gen-associated, or common if its sequence had significant sequence similarity (below the 10^-7^ e-value threshold) to genes found in the genomes of only bacterial pathogens, only non-pathogens, or both pathogens and non-pathogens, respectively. With the original intent of finding functional genes widely conserved among the diverse pathogens/non-pathogens, the unique genes were subdivided into “high quality” and “low quality” pathogen-associated genes or non-pathogen-associated genes based on three criteria: 1) broad phylogenetic distribution (conserved in three or more bacterial genera), 2) non-genus specificity (at least one genus in which the non-common gene was detected must contain both pathogenic and non-pathogenic species), and 3) proteins with probable function (more than 100 amino acids in length). Pathogen-associated genes and non-pathogen-associated genes that satisfied all three criteria are considered “high quality” while the others are considered “low quality.”

To refine the results from the pathogen-associated genes analysis, OrthoFinder v2.3.12 (15) was used to infer homologous relationship among all bacterial genes and cluster homologs into “orthogroups.” A species tree and gene trees of each orthogroup are then constructed for gene tree-species tree reconciliation to identify gene duplication/loss and to infer probable paralogs within each orthogroup. The NCBI RefSeq bacterial genome dataset was reduced to 501 reference and representative genomes prior to orthology inference by OrthoFinder(15). Pathogen-associated and non-pathogen-associated orthogroups were identified if all orthologs within the orthogroups belonged to genomes of pathogenic or non-pathogenic bacteria, respectively. Taxonomic conservation of pathogen-associated and non-pathogen-associated orthogroups were assessed by counting the number of unique bacterial genera from which their orthologs were detected.

To address the over-representation of certain species like *P. aeruginosa,* with many sequenced clinical isolates, within the original genome dataset, pairwise MASH distances were used to determine a genomic distance cut-off for clustering highly similar genomes and to select a single representative genome from each cluster. Specifically, the MASH distance matrix was used for hierarchical clustering and dendrogram visualization of all *Pseudomonas* genomes, using the R “stats” package v4.0.2. Upon the examination of a few tested MASH distance cut-offs of 0.21, 0.1, 0.08, 0.06 and 0.04 which generated 133, 206, 237, 295, and 380 genome clusters, respectively. Within each cluster under the 0.1 MASH (55) distance cut-off, an NCBI reference or representative genome was prioritized for selection. If no reference or representative genome was present in the cluster, one genome is randomly selected.

### Evolutionary selection inference

Evolutionary analysis of PAGs in 14 *P. aeruginosa* representative genomes was performed using the *HyPhy* v2.5, which incorporates synonymous rate variation into positive selection inference(20, 56). For each gene set, a codon-based multiple nucleotide sequence alignment, with masked internal and external stop codons, was generated from coding sequences for all *P. aeruginosa* genes within the orthogroup using MACSE v2.04 (57).

Multiple sequence alignments of the gene sets of interest are preprocessed and screened for recombination by HyPhy’s Genetic Algorithm for Recombination Detection (GARD) method (58). Genes were analyzed for positive selection by multiple HyPhy methods. Mixed Effects Model of Evolution (MEME) was chosen as the primary method for detecting site-specific episodic positive selection as it has more power than other methods in detecting individual sites that have evidence of positive selection under a proportion of the branches in a phylogeny (19, 59).

Adaptive Branch-Site Random Effects Likelihood (aBSREL) was used complementary to MEME for identifying specific branches on a phylogeny on which positive selection is evident(60). Contrary to the three aforementioned algorithms which test for episodic selection, the Fixed Effects Likelihood (FEL) was used to test for pervasive positive selection in small (less than 100 sequences) gene sets such as the pathogen-associated genes and T3SS-related genes dataset (20). Finally, the Branch-Site Unrestricted Statistical Test for Episodic Diversification (BUSTED) method, identified gene-wide instead of site-specific episodic positive selection (19, 61).

RNA polymerase sigma factor (*rpoD*), DNA gyrase subunit B (*gyrB*), and DNA-directed RNA polymerase β chain (*rpoB*) were used as negative controls, as purifying selection for these three genes has been supported by published data (62). Likewise, the pyoverdine outer membrane receptor (*fpvA*) and the pyoverdine inner membrane protein (*fpvG*), with published evidence of positive selection (63), were used as positive controls in this analysis.

### Structure-function prediction of PvmSR using AlphaFold and Foldseek

PvmS (PA14_31050) and PvmR (PA14_31060) were first functionally characterized through similarity searches in the non-redundant protein sequences database (nr) using the NCBI BLASTp tool with default settings (64). Next, both structural and sequence alignments of PvmSR to similar proteins and *S. aureus* BlaIR were conducted using FoldMason with the Foldseek search server (39). First, 3D structures of PvmS and PvmR were generated with AlphaFold 3 using the AlphaFold server (65). From the results of the structure predictions, the atypical BlaIR orthologs from *M. tuberculosis* and *M. abscessus* found in Bonefont *et al.* (28) were of interest, due to the absence of the cytoplasmic sensor domain in the BlaR proteins of both species – much like in PvmS. To identify any other orthologs, 3D structures of PvmSR were then searched against all databases in the Foldseek search server to identify any orthologs present in pathogenic species. Based on the search results, the following protein structures were then aligned in two FoldMason MSAs: (1) *P. aeruginosa* PA14 PvmS (PA14_31050), *S. aureus* NCTC 9789 BlaR1, *M. tuberculosis* H37Rv Rv1845c, *M. abscessus* ATCC 19977 MAB_2414c, *M. leprae* TN ML2064 and *N. brasiliensis* ATCC 700358 O3I_019585; and (2) *P. aeruginosa* PA14 PvmR (PA14_31060), *S. aureus* NCTC 9789 BlaI, *M. tuberculosis* H37Rv Rv1846c, *M. abscessus* ATCC 19977 MAB_2415c, *M. leprae* TN ML2063 and *N. brasiliensis* ATCC 700358 O3I_019590. MSAs generated by FoldMason were then visualized with Jalview (v2.11.5.1, (66)) and coloured by conservation, and the structural alignments were visualized in ChimeraX (v1.11.1)(67).

### Generating deletion mutants, complementation and whole-genome sequencing confirmation

#### Strains and media conditions

*E. coli* OP50 was cultivated on Lysogeny Broth (LB) Lennox conditions (5 g/L NaCl) in broth and 1.5% agar at 37°C. *P. aeruginosa* PA14 and its mutants were cultivated on LB Luria conditions (0.5 g/L NaCl) in broth and 1.5% agar at 37°C. Bristol N2 *C. elegans* were cultivated and maintained on NGM medium seeded with *E. coli* OP50 and grown at 20°C(68). Additional growth media and conditions are outlined in subsequent methods.

#### Knock-out mutant generation

The PA14_31050/31060 deletion mutant (“ΔpvmSR”) was generated utilizing homologous-recombination of the suicide vector pEX18Gm and subsequent counter selection of the *sacB* gene product (69). Genomic DNA was extracted from *P. aeruginosa* PA14 using the DNeasy Blood & Tissue Kit (Qiagen). Separate upstream and downstream 500 bp regions of *pvmR* were amplified using the respective primers (**Table S7**) (5’ – 3’); 31060F1BamH1 – cctggatccGCTACGACGCGCAGCGCATGG, 31060R1 – CTGAAGTAGTGGGCCATAGCTCGTCAGGCACCAGCGAGAAGCCCATGC, 31060F2 – CGGCATGGGCTTCTCGCTGGTGCCTGACGAGC-TATGGCCCACTACTTCAG, and 31060R2HindIII – ccgaagcttCAC-CAGCGAGGCAAGGGTTCGAC. The lower-case sequences were designed with the desired restriction sites for insertion into pEX18Gm. The amplicons were generated by Phusion polymerase (Thermo Fisher Scientific) and purified by the GeneJet Gel Extraction Kit (Thermo Fisher Scientific). The upstream fragment was digested with BamHI, the downstream amplicon was digested with HindIII and the vector pEX18Gm was digested with both. The digested fragments were cleaned with the GeneJet Gel Extraction Kit and ligated together with T4 Ligase (Invitrogen Life Technologies) at a ratio of 2:1 (insert:vector). The resulting reaction was transformed into TOP10 *E. coli* competent cells (Invitrogen Life Technologies) and plated onto LB gentamicin (GEN) 15 μg/mL agar plates. Transformants were cultured and their plasmids purified using the GeneJet Mini Prep Kit (Thermo Fisher Scientific); clones were confirmed via Sanger sequencing (UBC Sequencing and Bioinformatics Consortium). The purified plasmid was transformed into ST18 *E. coli* cells; plates supplemented with aminolaevulinic acid (ALA).

The resulting strain was cultured, diluted to an OD600 = 0.1 and mixed at a 2:1 ratio with PA14, and spotted onto LB plates with ALA. The cells were recovered in sterile LB and spread onto 50 μg/mL GEN LB plates. Single colonies were then patched onto 10% sucrose LB plates. The patching was repeated for 4 days and subsequently the sucrose was increased to 12%, and patching repeated two more times at the increased sucrose concentration. For the 6^th^ patch, the colonies were cultivated on both 12% sucrose LB and 50 μg/mL GEN LB plates. Colonies that grew on sucrose plates and were sensitive to GEN were sequenced to confirm the deletion of *pvmR*.

#### Genomic DNA extractions and Oxford Nanopore whole genome sequencing

Bacterial cultures were grown ∼18 h at 37°C in LB medium. 500 µL of culture were pelleted and genomic DNA purified using Qiagen DNeasy Blood & Tissue kit using the modifications for gram-negative bacteria. Sample quality was confirmed using Nanodrop (Thermofisher Scientific), DNA electrophoresis, and fluorescence quantitation (Biotium). 300 ng of genomic DNA was prepared for sequencing using the Rapid Barcoding Kit 96 V14 (SQK-RBK114.96) and loaded onto a FLO-MIN114 MinIONION flow cell (R10.4.1), with high-accuracy basecalling using MinKNOW Dorado (version 7.4.13). Genome assembly was performed using flye (version 2.9)(70).

#### *C. elegans* slow-killing virulence assays

Virulence determination was performed using *E. coli* OP50 and *P. aeruginosa* PA14 strains in an established virulence assay using the *C. elegans* infection model (29). *C. elegans* (wild-type Bristol N2) were synchronized by embryo isolation and maintained on nematode growth medium (NGM) plates coated with *E. coli* OP50 as food source (68). To prepare for the slow-killing assay, low-osmolarity NGM plates were seeded with 50 μL of overnight culture of the *P. aeruginosa* strains as noted, or *E. coli* OP50 as positive control, per 3.5 cm diameter plates for bacterial lawn formation at 37°C overnight. The plates were then equilibrated at 20°C overnight prior to infection. To prevent progeny development during the assay, 100 mM FUdR was added to the NGM plates to a final concentration of 50 or 75 μM as noted. 30 L4 stage worms were transferred onto each plate, incubated at 20°C and were monitored for viability for up to 8 days (196 hours). Worms are considered dead if they no longer moved or responded to touch. The slow killing assay was performed for a minimum of three biological replicates, each with three technical replicates.

Kaplan-Meier survival curves were estimated and visualized for each set of worms grown in the presence of 1) *E. coli* OP50 (negative control), 2) wildtype *P. aeruginosa* PA14 (positive control) and 3) the deletion mutant *ΔpvmSR*. Differences in worm survival between infection with the wildtype and mutant strains of *P. aeruginosa* were assessed using a log-rank test with a statistically significant p-value threshold of 0.05. Survival curve analysis and visualization were done with the R packages “survminer” v0.4.8.

#### In vitro characterization of deletion mutants in β-lactam resistance and biofilm formation Minimum Inhibitory Concentration (MIC) Broth Microdilution

The protocol for minimum inhibitory concentration (MIC) experiments was adapted from Wiegand *et al.* 2008. Briefly, overnight cultures were grown in triplicate in cation adjusted Mueller Hinton Broth (MHB) or LB as noted. In a 96-well untreated, round bottom polystyrene plate, sterile media was added to the exterior rows and columns of the plate to prevent evaporation and edge effects. In the interior wells, serial dilutions of the appropriate antibiotics were performed in 50 μL volumes. Cultures were then diluted to a final OD600 of 0.001 (∼5 x 10^5^ – 5 x 10^6^ colony-forming units (CFU) mL^-1^), and 50 μL of diluted bacteria was added to each interior well. The MIC plate was incubated overnight and, the following day, growth was recorded visually when a cell mass at the bottom of the well was visible. Growth was also measured with a plate reader using OD600 values.

#### *C. elegans* slow-killing virulence for Dual-RNASeq

The experimental set-up for the Dual-RNASeq experiment follows the outlined slow-killing virulence assay above with the following modifications. 100 L4 stage worms were moved to the experimental plates instead of 30. Following incubation at 20°C for 24 h, the worms were recovered from the plates with 1X M9 minimal media and washed 3 times. Following the final wash, 1 mL of TRIzol was added to the tube, the worms suspended and frozen at –80°C. After the experiment was performed in triplicate, the samples were processed simultaneously. The samples were thawed at 37°C and an additional 2 mL of TRIzol were added to the samples. The samples were then subjected to six additional freeze/thaw cycles. When most worms integrity appears disrupted under a dissecting scope, 200 μL of chloroform was added to the tubes, inverted for 15 s and allowed to separate into phases for 3 min before spinning at 12,000 x g for 15 min at 4°C. RNA in the aqueous layer was then further purified using an RNeasy Mini Kit (Qiagen) following the protocol for Purification of Total RNA from Animal Cells using the optional on column DNA digest with DNase I (ThermoFisher Scientific). The eluted RNA was checked for quality and integrity using Nanodrop (ThermoFisher Scientific) and Bioanalyzer (Agilent Technologies). The RNA was prepared for sequencing using the Illumina Stranded Total RNA Prep with Ligation, Ribo-Zero Plus Microbiome and the library quality and integrity was confirmed by TapeStation (Agilent Technologies). The samples were sequenced at the Genome Sciences Centre (Vancouver, BC, Canada); the library was initially sequenced on a MiSeq (Illumina) for quality control, before PE150 sequencing on an Illumina NovaSeq.

#### Dual-RNASeq analyses to assess transcriptional changes in both host and pathogen

Raw sequence reads were checked for sequencing quality (FastQC v0.12.1) and aligned against a genome constructed by concatenating *C. elegans* and PA14 reference genomes (STARv2.7.11b (71); reference genomes: GCF_000002985.6_WBcel235 from NCBI on January 21, 2026, UCBPP-PA14_109 from pseudomonas.com on January 27, 2026). Due to the *pvmR* (PA14_31060) annotation found only in the GFF v3 for PA14, the reference genome annotation was constructed by first converting the PA14 GFF v3 annotation to GTF v2.2 using AGAT v1.6.1 before concatenating with the GTF v2.2 annotation for *C. elegans*. Count tables were generated using htseq-count v2.0.5(72); uniquely mapped reads were separated into *C. elegans* and PA14 annotation identifiers unique to each organism. The number of uniquely mapped reads ranged from 1.2M - 3.7M (median of 2.4M) for *C. elegans* and 1.1M - 5.1M (median of 2.4M) for *P. aeruginosa*. DESeq2 v1.52.0 (73) was used to determine DEGs with an adjusted p value of ≤ 0.05. Functional enrichment of DEGs from *C. elegans* were performed using KEGG pathway annotations (74) through clusterProfiler v4.20.0 (75) (reference organism *C. elegans* (nematode), and Wormcat v2.0.1 (76) using the provided “whole_genome_v2_nov-11-2021.csv” annotation file from its R package. Functional enrichment of DEGs from PA14 were analysed for over-representation analysis of GO terms using annotations from eggnog-mapper v2.1.13 (77) on the database v5.0.2 with “--sensmode ultra-sensitive” for Diamond, and “--go_evidence All” options from Galaxy Europe before calculating enrichment through clusterProfiler. Significantly enriched GO terms and KEGG pathways were assessed within clusterProfiler using a hypergeometric distribution with multiple test correction using the Benjamini-Hochberg method for an adjusted p value ≤ 0.05. Wormcat uses Fisher’s exact tests and Bonferroni multiple test correction to calculate significance, which were similarly filtered for an adjusted p value of ≤ 0.05. Volcano plots were created using EnhancedVolcano v1.30.3 (https://github.com/kevinblighe/EnhancedVol-cano#references), gene-concept networks were created using enrichplot clusterProfiler, heatmaps were created pheatmap v1.0.13 (https://cran.r-universe.dev/pheatmap/citation) and any remaining plots were created using ggplot2 v4.0.3 (https://ggplot2.tidyverse.org/) in R v4.6.0 (https://www.r-project.org/).

## Data availability

All data have been deposited in Gene Expression Omnibus (GEO).

